# An Epiblast Stem Cell derived multipotent progenitor population for axial extension

**DOI:** 10.1101/242461

**Authors:** Shlomit Edri, Penny Hayward, Peter Baillie-Johnson, Benjamin Steventon, Alfonso Martinez Arias

**Affiliations:** Department of Genetics Downing Site University of Cambridge Cambridge, CB2 3EH UK

## Abstract

The Caudal Lateral Epiblast of mammalian embryos harbours bipotent progenitors that contribute to the spinal cord and the paraxial mesoderm in concert with the elongation of the body axis. These progenitors, called Neural Mesodermal Progenitors (NMPs) are identified as cells coexpressing *Sox2* and *T/Brachyury*, a criterion used to derive NMP-like cells from Embryonic Stem Cells in vitro. However, these progenitors do not self renew, as embryonic NMPs do. Here we find that protocols that yield NMP-like cells in vitro first produce a multipotent population that, additional to NMPs, generates progenitors for the lateral plate and intermediate mesoderm. We show that Epiblast Stem Cells (EpiSCs) are an effective source for these multipotent progenitors that are further differentiated by a balance between BMP and Nodal signalling. Importantly, we show that NMP-like cells derived from EpiSCs self renew in vitro and exhibit a gene expression signature similar to that of their embryo counterparts.

## Introduction

The anteroposterior axis of a vertebrate can be subdivided into three anatomically distinct regions: the head, the trunk and the tail. The trunk starts at the end of the hindbrain, runs to the anus and comprises derivatives of the mesoderm and the ectoderm such as the thoracic cage, muscles, kidneys and spinal cord. The thoracic tract has different origins in different organisms: in anamniotes e.g. fish and frogs, it is inferred to arise during gastrulation from a pool of pre-existing cells within the ectoderm, while in amniotes e.g. chickens and mice, it is derived from the expansion of the Caudal Epiblast (CE), a proliferative region located at the caudal end of the embryo, where the primitive streak persists, that acts as a source for paraxial, intermediate and lateral plate mesoderm as well as for the spinal cord (Henrique et al., 2015; Stern, 2005; Steventon and Martinez Arias, 2017; Sweetman et al., 2008; Wilson et al., 2009). Lineage tracing studies have shown that the CE harbours a population of bipotential progenitors located behind the node, at the Node Streak Border (NSB) and extending laterally into the Caudal Lateral Epiblast (CLE), that give rise to neural and mesodermal precursors. These cells have been called Neural Mesodermal Progenitors (NMPs), are often characterized by simultaneous expression of *T* (also known as *Bra*) and *Sox2* (Cambray and Wilson, 2007; Wymeersch et al., 2016) and are capable of limited self renewal (Cambray and Wilson, 2002; McGrew et al., 2008; Tzouanacou et al., 2009).

Recently a few studies have claimed the generation of NMP-like cells in adherent cultures of mouse and human embryonic Pluripotent Stem Cells (PSCs) (Gouti et al., 2014; Lippmann et al., 2015; Turner et al., 2014). In these studies, Embryonic Stem Cells (ESCs) are coaxed into a transient *T* and *Sox2* coexpressing state that, depending on the culture conditions, can be differentiated into either paraxial mesoderm or spinal cord progenitors and their derivatives. However, there is no evidence that these NMP-like cells are propagated in vitro as they are in the embryo (Tsakiridis and Wilson, 2015). Furthermore, coexpression of *T* and *Sox2* might not be a unique characteristic of NMPs as it is also a signature of Epiblast Stem Cells (EpiSCs) (Kojima et al., 2014) which are pluripotent and this does not imply that EpiSCs are NMPs. While other markers have been used to refine the molecular identity of NMPs in vitro e.g. *Nkx1-2, Cdx2, Cdh1* and *Oct4*, these are also expressed in the epiblast and in the primitive streak during gastrulation (see Supplementary section 1-2 and Fig. S1), emphasizing the notion that these gene expression signatures are not uniquely associated with NMPs. Altogether these observations raise questions about the identity of the *T* - *Sox2* coexpressing cells derived from ESCs and about the signature of the NMPs.

Here, we show that *T* - *Sox2* coexpressing cells derived from ESCs and EpiSCs based differentiation protocols display differences at the level of gene expression and represent different developmental stages of the transition between naïve, primed pluripotency and neuro-mesodermal fate choices. Furthermore, we find that, in adherent culture, all available protocols generate a multipotent population where in addition to an NMP signature we find also signatures for Lateral Plate and Intermediate Mesoderm (LPM and IM) as well as the allantois. We report a new protocol, based on EpiSCs, that sequentially generates the multipotent population and an NMP-like population with many of the attributes of the embryonic NMPs. In particular these cells exhibit an ability to self renew in vitro and to contribute to posterior neural and mesodermal regions of the embryonic body in a xenotransplant assays. Our study leads us to propose that, in vitro and in vivo, NMPs are derived from a multipotent population that emerges in the epiblast at the end of gastrulation and gives rise not only to the elements of the spinal cord and paraxial mesoderm but all elements of the trunk mesoderm. Our study also provides a culture system for the establishment of a self renewing NMP-like population in vitro.

## Results

### EpiSCs yield a postimplantation epiblast population that resembles the CLE

Several protocols allow the differentiation of ESCs into an NMP-like population, defined as cells that coexpress *T* and *Sox2*, that can be further differentiated into neural and mesodermal progenitors (summarized in (Henrique et al., 2015)). However, it is not clear whether these NMP-like cells derived through different protocols are similar to each other and, importantly, how each relates to the NMPs in the embryo. To begin to answer these questions we compared NMP-like cells obtained from three different protocols: ES-NMPs (Turner et al., 2014) and ES-NMPFs (Gouti et al., 2014), derived from ESCs, as well as Epi-NMPs, derived from a new protocol that we have developed from EpiSCs (Fig. 1a; see Methods). All protocols yield cells coexpressing *T* and *Sox2* at the level of both the mRNA and protein (Fig. 1b and Supplementary Fig. S2), but differ in the levels and degree of correlated expression of the two genes (Fig. 1b and Supplementary Fig. S2b) and the Epi-NMP population has a highest number of *T* - *Sox2* positive cells in a transcriptional stable state (as defined by a low variance of *T* and *Sox2*) in comparison to the other conditions (Supplementary Fig. S2b).

**Fig 1.**
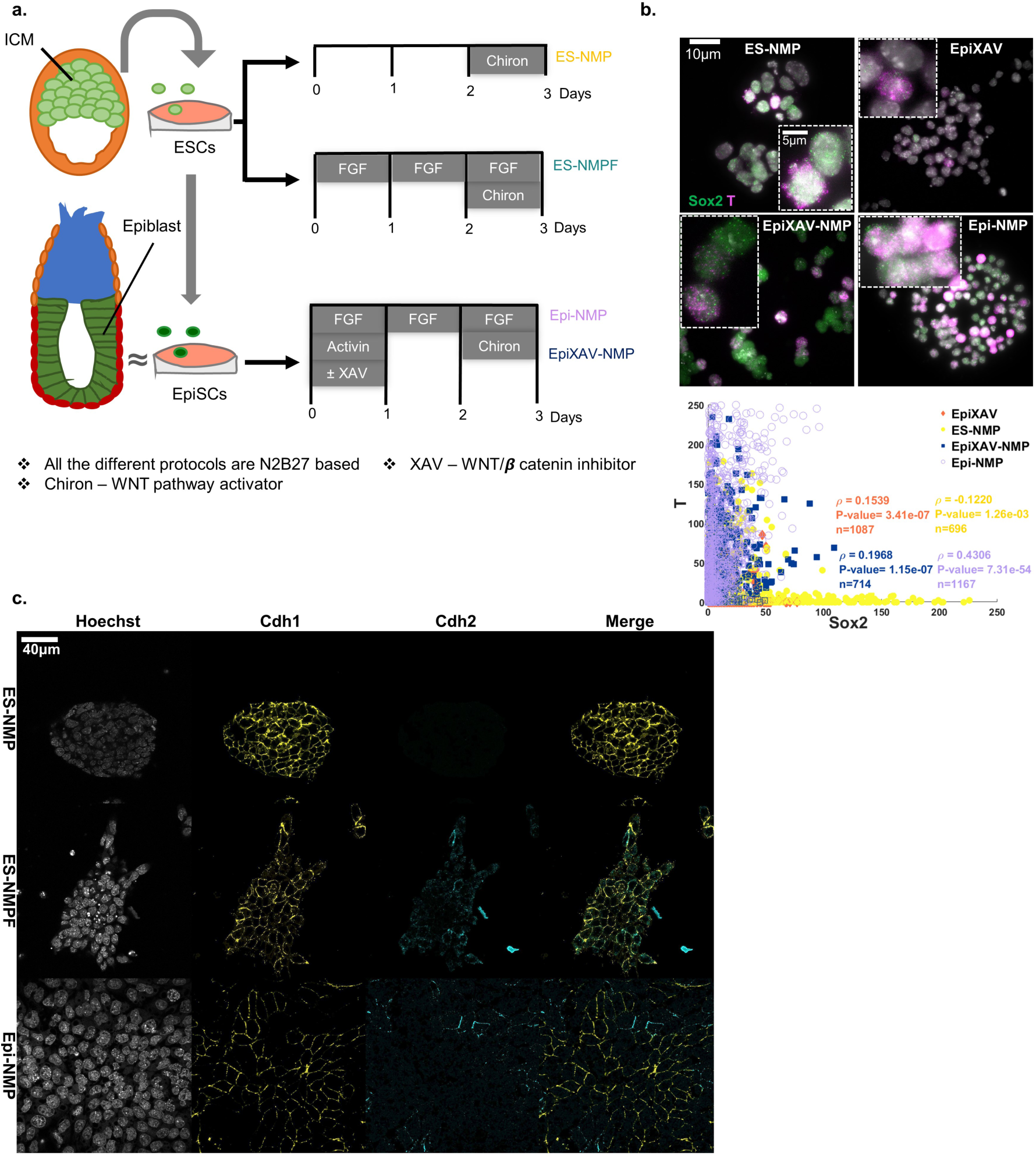
Comparison between in vitro protocols to produce NMP-like cells. **a**. Diagram of the protocols: ES-NMP (Turner et al., 2014) ES-NMPF (Gouti et al., 2014) and Epi-NMP (see Methods). **b**. images, obtained by using single molecule fluorescence in-situ hybridization (sm-FISH), of cells expressing *Sox2* (in green) and *T* (in magenta) mRNA in different conditions: ES-NMP, EpiXAV, EpiXAV-NMP and Epi-NMP. The insets are zoom in on cells coexpressing *Sox2* and *T*. Quantification plot of the number of mRNA molecules in a cell in the different conditions can be found underneath the images. Each dot represents a cell where the x-axis and y-axis represent the number of *Sox2* and *T* molecules respectively in a cell. The Spearman coefficient correlation between *Sox2* and *T*, the P-value and the total number of cells, noted as ρ, P-value and n respectively, can be found as an inset in the quantification plot. **c**. Confocal images of ES-NMP, ES-NMPF and Epi-NMP on their 3^rd^ day. Nuclei (Hoechst in grey), *Cdh1* (yellow) and *Cdh2* (cyan) (see Methods).

To characterize the different NMP-like populations further, we investigated the expression of several genes associated with the epiblast, the CE and the NSB/CLE region, where NMPs are thought to reside, between stages E7.0 and E8.5 (see Supplementary sections 1-2 and Fig. S1 for the criteria we followed to select these genes). Both of the ES derived NMP-like populations exhibit clear expression of *Cdh1* and *Oct4* and low levels of *Fgf5* and *Otx2*, with ES-NMPF cells specifically displaying high levels of genes associated with mesendoderm differentiation e.g. *Mixl1* (endoderm), *Tbx6* (paraxial mesoderm) and *Evx1* (extraembryonic mesoderm) (Fig. 2b, Supplementary Fig. S4 and see also (Gouti et al., 2014)). This suggests that ES-NMP and ES-NMPF are overlapping populations with an overrepresentation of cells in the early epiblast/gastrula like stage. In contrast, Epi-NMPs are in a very different state: in addition to T, *Sox2* and *Nkx1-2* these cells express significant levels of *Nodal, Fgf8, Fgf5, Foxa2, Otx2* and *Oct4* together with *Cyp26a1* (see Fig. 2b and Supplementary Fig. S4). This is a profile associated with the late epiblast (about E7.5), around the time of the appearance of the CE, before NMPs can be detected (see Supplementary sections 1-2 and Fig. S1).

**Fig 2.**
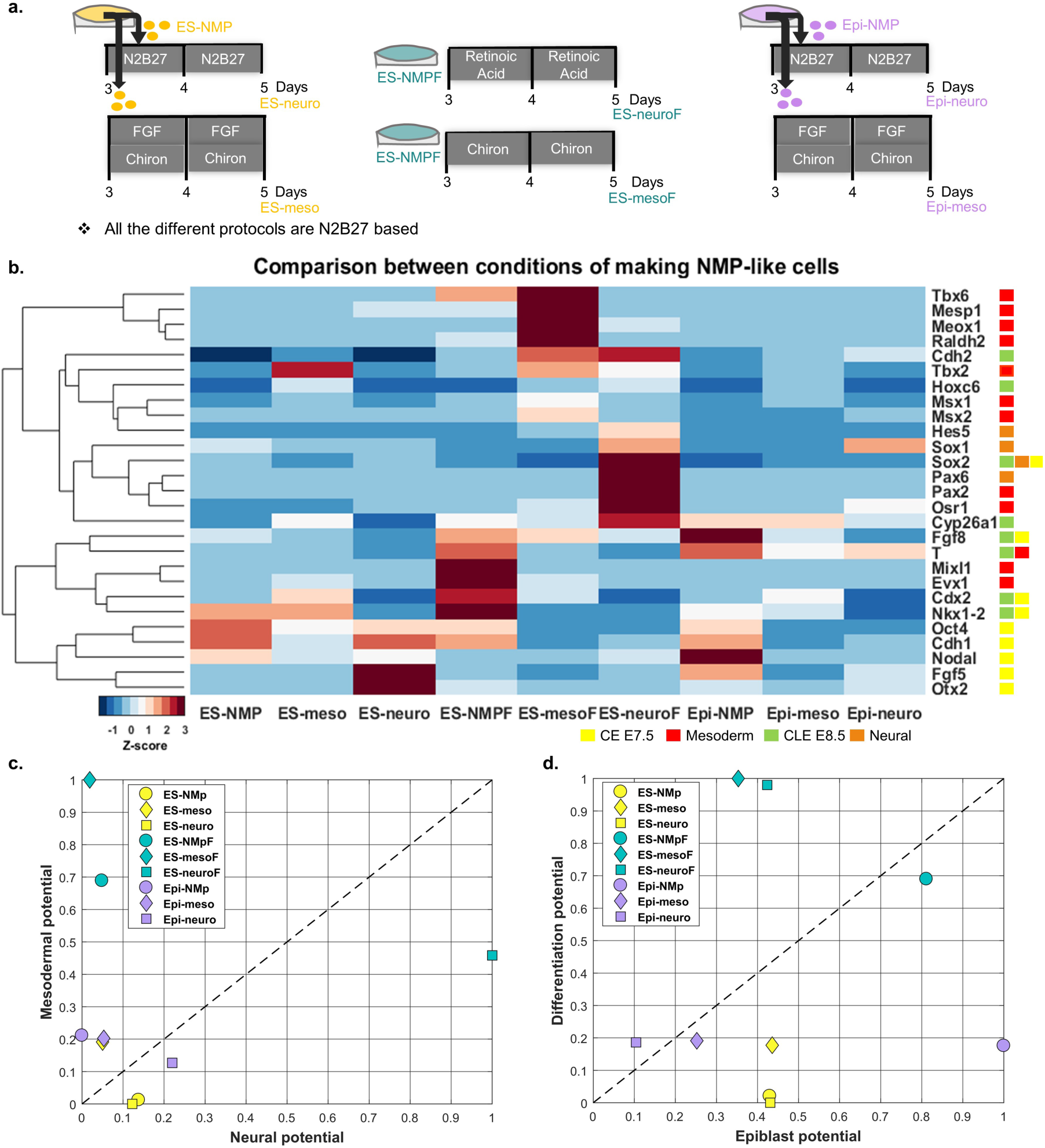
Differentiation NMP-like cells into neural and mesodermal precursors. **a**. ESCs and EpiSCs were grown for 3 days accordingly to specific protocols (Fig. 1a; see Methods). After 3 days, the ES-NMP (yellow) and the Epi-NMP (purple) were split into 2 flasks and cultured for 2 days in a medium that allows differentiation to either neural or mesodermal cells (see Methods). In the case of the ES-NMPF (turquoise) we followed the published protocol (Gouti et al., 2014) and did not split/passage the cells, which were grown for 5 days in the same flask in the neural or mesodermal conditions (see Methods). We named the resulting populations ES-neuro/ES-neuroF and ES-meso/ES-mesoF for those with an ES-NMP/ES-NMPF origin, and Epi-neuro and Epi-meso for those with an Epi-NMP origin. **b**. Expression heatmap of 27 genes, obtained by RT-qPCR, in cells grown in the different conditions, as indicated in Fig. 2a. The expression of each gene was normalized to the expression in the Epi-meso condition and then was scaled across the different conditions via calculating the Z-score (see Methods and Supplementary Fig. S4). Each gene was assigned to a label according to Fig. S1 and Supplementary sections 1-2: CE E7.5 in yellow, mesoderm in red, CLE E8.5 in green and neural in orange. **c**. calculation of the NMP index. In all the conditions the average expression Z-score value of the mesodermal genes (marked in red) and the neural genes (marked in orange) were calculated and scaled between 0 – 1 across conditions. Those are the Neural potential in the x-axis and Mesodermal potential in the y-axis (see Methods), highlighting the neural-mesodermal state of each condition. **d**. Calculation of the epiblast index. In all the conditions the average expression Z-score value of the differentiating genes (marked in red and orange) and the epiblast genes (marked in yellow and green) were calculated and scaled between 0 − 1 across conditions. Those are the Epiblast potential in the x-axis and Differentiation potential in the y-axis (see Methods), highlighting the epiblast state of each condition.

To better comprehend the gene expression profiles of the different culture conditions we calculated two measures of the degree of differentiation of each population: the ‘NMP index’ as a measure of the neuro-mesodermal bias of each population, and the ‘Epiblast index’, as a measure of the degree of differentiation of the population (Fig. 2c-d see Methods). In both cases, the distance of the cells from the diagonal and the origin reflects the average phenotype of the cell population: the closer to both the more the population is in a progenitor, uncommitted epiblast state. These indexes show that Epi-NMPs have the highest epiblast potential with a slight differentiation bias towards the mesoderm, while ES-NMPs exhibit low epiblast potential with a degree of differentiation towards the neural fate. In contrast, the ES-NMPFs exhibit high epiblast and mesodermal potential. The differences between the three populations are further emphasized by an examination of the protein levels of some of these markers (Fig. 1c, Supplementary Fig. S2a and Fig. S3). NMP-like populations derived from ESCs exhibit high levels of *Sox2, Oct4* and *Cdh1* expression with some cells expressing *Otx2*, a signature characteristic of early epiblast (Morgani et al., 2018). In the case of ES-NMP-like there is no expression of *Cdh2*, whereas for ES-NMPF we observe a combination of *Cdh1* and *Cdh2* at the level of single cells, a situation rarely seen in vivo (Fig. 1c). In contrast, the Epi-NMP exhibit lower level of *Sox2* and *Oct4* (Supplementary Fig. S2a) and a mutual exclusive expression of *Cdh1* and *Cdh2* (Fig. 1c), which is a characteristic of the late Epiblast (Corsinotti et al., 2017; Morgani et al., 2018).

Exposure of the different NMP-like populations to neural and mesodermal differentiation environments reveals their potential (Fig. 2 and Fig. S4; see Methods). In all cases the cells differentiated into neural and mesodermal progenitors but exhibited biases depending on their origin (Fig. 2b-d): ES-NMPFs and its differentiated progeny exhibit a mesodermal bias while ES-NMPs exhibit a bias towards the neural fate. In contrast, Epi-NMPs do not exhibit any strong bias.

Altogether these results suggest that different protocols yield related but different NMP-like populations which might have different functional properties. The NMP-like population derived from EpiSCs, being the closest to the epiblast and its differentiated progeny to the NMP potential.

### Developmental staging of in vitro derived NMP populations

The differences between the candidate NMP-like populations derived in vitro suggest that they might represent different stages of the transition between the early postimplantation epiblast and the CLE. To test this, we created a developmental stage reference using a microarray study of the epiblast at different embryonic stages between early postimplantation (E5.5) and early-CLE (E8.25) (Kojima et al., 2014), and mapped the NMP-like populations, as well as their differentiated derivatives, onto it (Supplementary section 3 and Fig. S5). Using this as a reference, we observe that in a three-dimensional Principal Component Analysis space, ES-NMP, Epi-NMP and their derivatives mapped closely to the different embryonic stages, whereas the ES-NMPF and its neural and mesodermal differentiation populations are separate from these conditions and from the embryonic stages (Supplementary Fig. S5c). The Epi-NMP and its derivatives projected closely to each other within the embryo trajectory between the LMS and EB stages.

We also used our developmental reference to explore the proximity of the in vitro derived populations to specific epiblast states in vivo. To do this, we used the microarray epiblast analysis of Kojima et al. 2014 as a reference and calculated the cosine similarity between the in vitro population and the different stages of the embryo as a metric for the proximity of each in vitro population to a particular embryonic stage (Fig. 3a and Supplementary Fig. S5b (see Methods). Using this measure we find that ES-NMPs correlate with early epiblast, whereas ES-NMPFs appear to be a broad population associated with different stages, mostly late epiblast, confirming that both represent heterogeneous populations of differentiating cells. On the other hand, the Epi-NMP population does not appear to show a clear similarity to a specific stage, other than to early epiblast and LS stages. This analysis also reveals that Epi-meso, a population derived from the Epi-NMPs, resembles the LB stages, which correspond to E8.25, the stage where the CLE can be seen to harbour the NMP for the first time.

**Fig 3.**
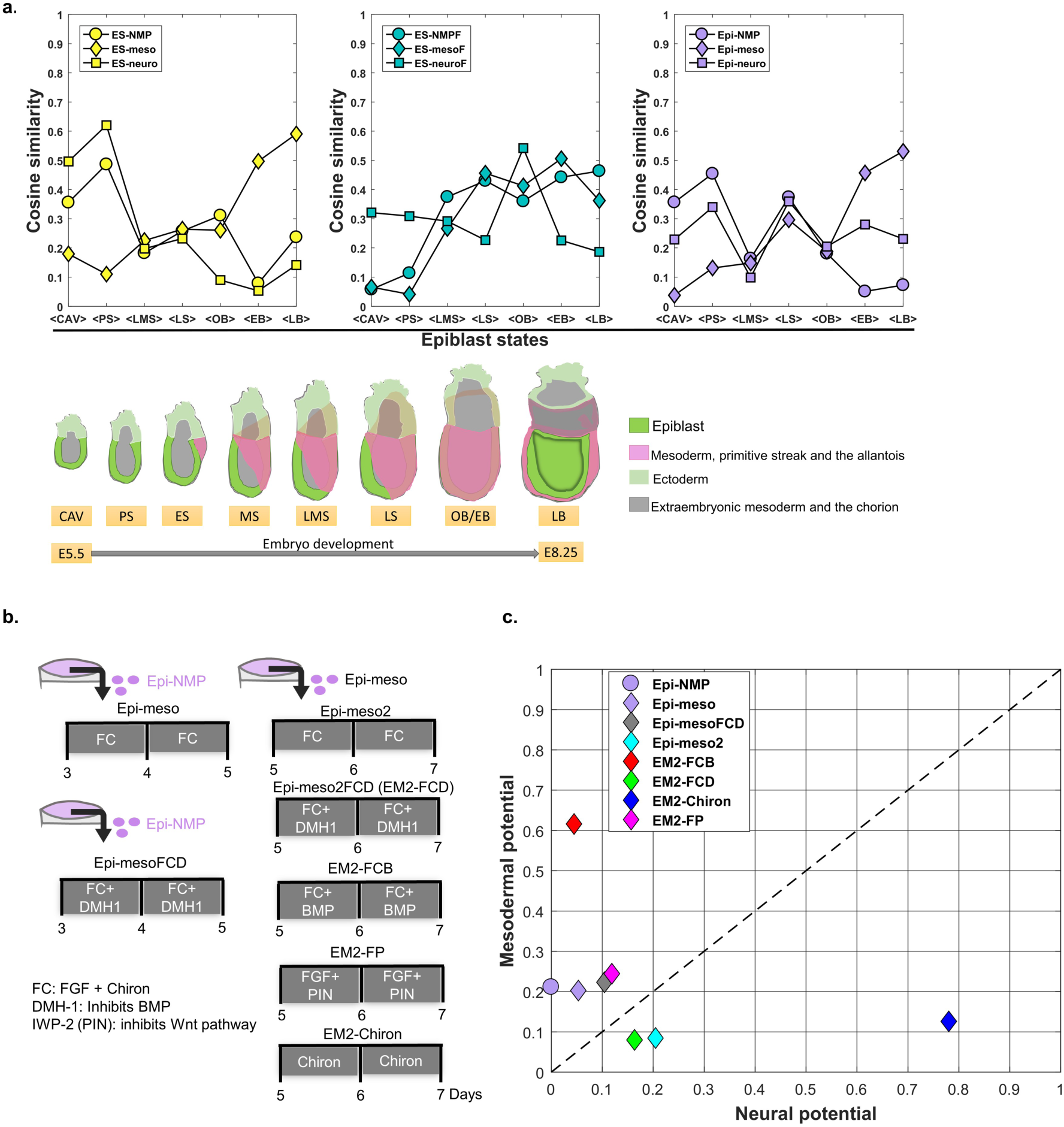
Comparison of the in vitro protocols to different epiblast stages and the effect of Wnt, FGF and BMP on mesodermal differentiation of the Epi-NMP population. **a**. Microarray gene expression data of the epiblast/ectoderm (excluding the primitive streak) from different stages of the mouse embryo (Kojima et al., 2014) was used as an anchor to compare the different protocols to the embryo; staging is shown underneath the similarity plots (after Kojima et al. 2014), where green indicates the epibast/ectoderm; pink the mesoderm, the primitive streak, and the allantois and grey the extraembryonic mesoderm and the chorion: CAV, cavity; PS, prestreak; ES, early-streak; MS, mid-streak; LMS, late mid-streak; LS, late streak; OB/EB, no bud/early bud; LB, late bud. The pairwise cosine similarity measure was calculated based on the expression of the 27 genes shown in Fig. 2b between the NMP in vitro protocols or their differentiation and the different stages of the epiblast mouse embryo (see Supplement Fig. S5a and Methods). The y-axis represents the average cosine similarity across the same epiblast mouse stages (x-axis) as shown in Fig. S5b. Value of 0 indicates dissimilarity and value of 1 indicates maximal similarity (see Supplement Fig. S5b and Methods). **b**. Differentiation protocols for Epi-NMP into Epi-meso and Epi-meso2 with modulation of BMP signalling (DMH-1 as an inhibitor and *Bmp4* as an agonist), Wnt signalling (IWP-2 an inhibitor of Wnt secretion as a Wnt signalling antagonist) and without the addition of exogenous FGF (see Methods). **c**. NMP index of the Epi-NMP and its differentiation protocols was calculated based on the Z-score of the normalized expression profile obtained by RT-qPCR shown in Supplementary Fig. S6c.

Altogether these results support the notion that different starting conditions and differentiation protocols lead to populations with different identities and representations: ES-NMP represents a heterogeneous early population, while ES-NMPF is a heterogeneous population dispersed over several stages with much differentiation. Epi-NMPs, on the other hand, appears to be a tighter population resembling a later epiblast.

### Multiple tail bud fates emerge from differentiating ESCs and EpiSCs in culture

In the course of our survey of cell type markers in the different populations, we noticed that all protocols that lead to coexpression of *T* and *Sox2* also lead to the expression of genes not associated with NMPs e.g. *Mesp1, Evx1, Mixl1, Gata6, Bmp4, Msx1, Msx2, Osr1, Pax2* and *Tbx2* (Fig. 2b and Supplementary Fig. S4 and see also (Amin et al., 2016; Gouti et al., 2014)). A survey of the literature shows that in the embryo between E7.0 and E8.5, roughly the stage of the differentiating in vitro cells, these genes are expressed in the posterior domain of the CE, in the progenitors of the allantois *(Tbx2, Tbx4, Mixl1* and *Evx1)*, the LPM *(Msx1* and *Msx2)* and the IM *(Pax2, Osr1)* (For further details see supplementary sections 1-2 and Fig. S1). This suggests that the three NMPs-like populations derived in vitro are not restricted to harbour NMPs only, but rather that they represent a multi-potential population which includes progenitors of LPM, IM and allantois.

In the embryo, the differentiation of the CE is under the control of BMP signalling, that favours more posterior fates (LPM, IM and allantois progenitors) at the expense of more anterior ones (NMPs) (Wymeersch et al., 2016). To test this, we altered the levels of BMP in the Epi-meso population, which appears to be the closest to the source of NMPs in the embryo (Fig. 3b-c, Supplementary Fig. S6). In our cultures, inhibition of BMP signalling elevates the expression of NMP markers e.g *T*, *Sox2* and *Cdx2* (Epi-meso versus Epi-mesoFCD in Supplementary Fig. S6c) and increases its NMP potential (Fig. 3c). On the other hand, addition of BMP to the derivatives of Epi-meso population (EM2-FCB) elevates dramatically their mesodermal potential (Fig. 3c) and specifically increases the levels of genes associated with posterior fates: *Bmp4, Msx1, Msx2* and *Tbx2* together with *Cdx2* and *Snail*, (Supplementary Fig. S6a, c). Similarly, to Epi-mesoFCD, inhibition of BMP in the Epi-meso2 population sample (EM2-FCD Fig. 3b-c), slightly improves its NMP index in comparison to Epi-meso2.

In the embryo, as the posterior region of the CE is dominated by BMP signalling, the differentiation of the anterior domain is dependent on its proximity to the NSB and we observe that the Epi-NMP population expresses *Nodal* and *Foxa2* genes which are associated with this region (see Fig. 2b and Supplementary Fig. S4a). Inhibition of Nodal signalling reduces *T -Sox2* coexpressing cells (Turner et al., 2014) and this led us to test whether Nodal signalling influence the NMP signature of this population. To do this, we cultured Epi-NMP from Nodal mutant EpiSCs (Nodal -/- Epi-NMP see Methods and Fig. 4a) and compared them to Nodal mutant Epi-NMPs supplemented with 2 different doses of Nodal in the presence of FGF and Chiron: 100ng/ml of Nodal (Nodal-/- Epi-NMP+0.1xNodal) or 1μg/ml of Nodal (Nodal-/- Epi-NMP+1xNodal). Loss of Nodal leads to an increment of *Sox2* and a decrease of *Cyp26a1* and *Fgf8* expression, suggesting a loss of CLE character, while addition of Nodal to these cells lifts their levels of *T* and lowers the levels of Sox2 expression. These results suggest that Nodal signalling is necessary to maintain the relative levels of *Sox2* and *T* and significant levels of *Fgf8* and *Cyp26a1*, which are characteristic of the CE.

**Fig 4.**
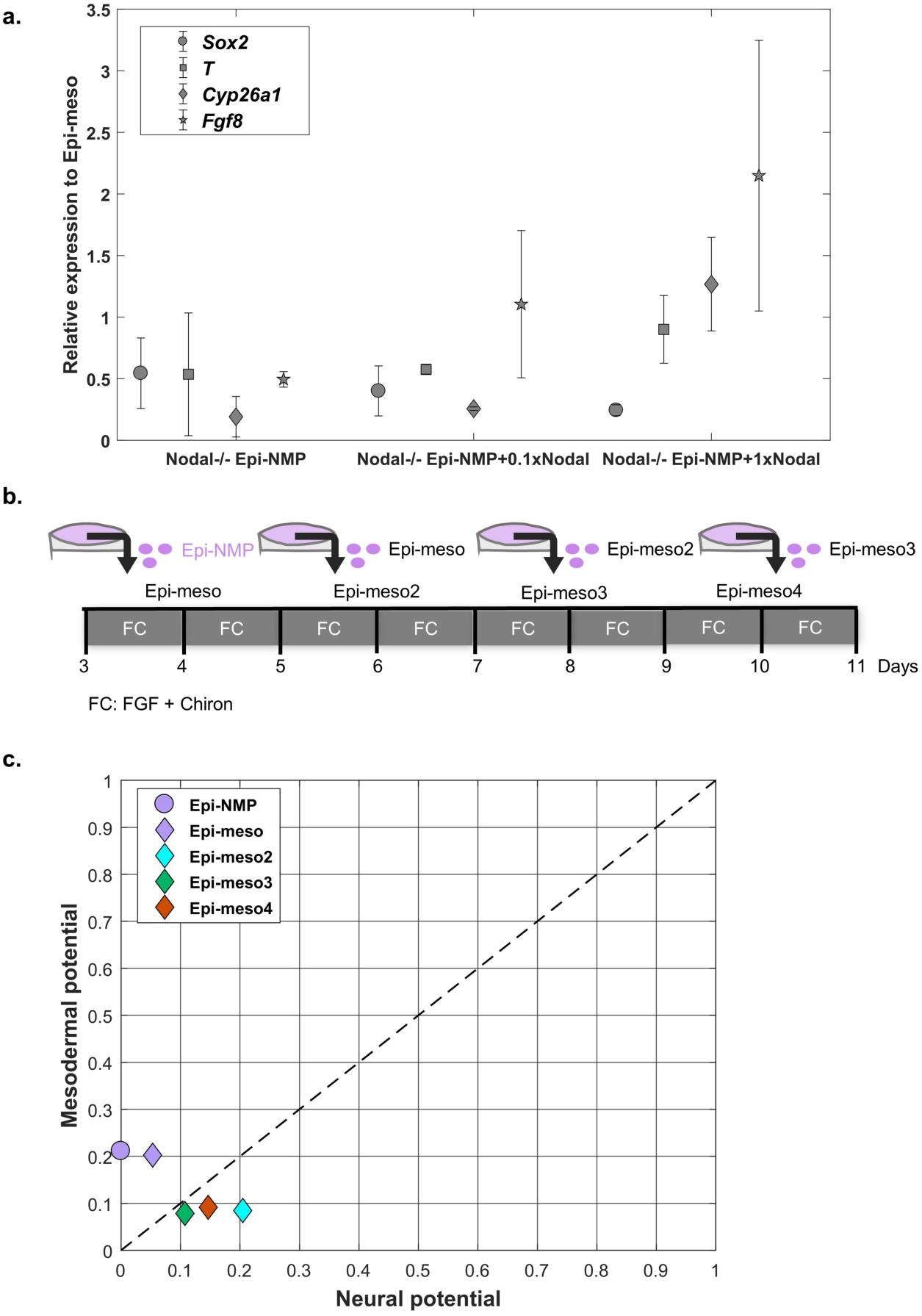
The effect of Nodal signalling and the maintenance of Epi-NMP in culture. **a**. Nodal mutant cells were cultured in the Epi-NMP protocol (Nodal -/- Epi-NMP). This population of cells was compared to the same population just with the addition of 2 doses of Nodal concentration to the growing medium of the Epi-NMPs on their 3^rd^ day: FGF, Chiron and either 100ng/ml of Nodal (Nodal-/-Epi-NMP+0.1xNodal) or 1μg/ml of Nodal (Nodal-/- Epi-NMP+1xNodal). The different shaped dots represent the genes average expression across biological replicas obtained by RT-qPCR and the error bars indicate the standard error between those replicas. The gene expression across the different conditions was normalized to Epi-meso condition (see Methods). **b**. Differentiation protocol for Epi-NMP into mesodermal precursors; cells were passaged and cultured in FGF and Chiron at every passage to generate the different generation of Epi-meso cells (Epi-meso, Epi-meso2, Epi-meso3, etc. see Methods). **c**. The NMP index of the Epi-NMP and its derivatives was calculated based on the Z-score of the normalized expression profile obtained by RT-qPCR shown in Supplementary Fig. S7b.

In summary, our results suggest that in all cases, differentiation of PSCs into a caudal population does not result in the specification of NMPs only, but rather of a multipotent population for all axial derivatives. This population is further differentiated by BMP and Nodal. The differences between the different protocols might not only result in different stages of development but also in different proportions of the different mesodermal populations.

### Epi-NMPs create a population that can be propagated in vitro

In the embryo, the initial NMP population needs to be amplified, together with the progenitors of the LPM and IM, if it is to account for the cellular mass along the length of the region posterior to the brain (Steventon and Martinez Arias, 2017; Wymeersch et al., 2016) and this amplification should be a criterion to identify NMPs in vitro. Earlier studies have shown that ESCs derived NMPs are not able to maintain the *T* - *Sox2* coexpressing cells when they are passaged in the conditions in which they were generated, namely FGF and Chiron or Chiron alone ((Gouti et al., 2014; Turner et al., 2014) and unpublished observations). Surprisingly, when Epi-NMPs are induced to differentiate into mesoderm by exposure to FGF and Chiron, they maintain *T* and *Sox2* expression for at least two passages (Epi-meso, Epi-meso2, Epi-meso3 (EM1, EM2, EM3; see Fig. 4b-c and Supplementary Fig. S7) with a low differentiation index. Furthermore, in the transition from Epi-NMP to Epi-meso, cells lose expression of epiblast markers e.g. *Fgf5, Nodal, Otx2, Oct4* and *Cdh1* (see Fig. 2b, Fig. S4a and protein expression of *Oct4, Otx2 Cdh1* and *Cdh2* in Fig. 1c and Supplementary Fig. S2a and Fig.S3), hence this suggest that Epi-meso population is a refined state containing many features of the NMPs that are a subset of the CLE.

During the passages of the Epi-meso populations, we observe a progressive decrease in the expression of NMP markers (*T*, *Cyp26a1, Fgf8* and *Nkx1-2)* and a slow increase in the expression of differentiation genes associated with neural fates: *Cdh2, Sox2*, and *Hes5* (Supplementary Fig. S7). When Epi-meso cells are grown in N2B27 supplemented with Chiron alone (EM2-Chiron, Fig. 3b-c and Supplementary Fig. S6b-c), we observe an increase in the levels of expression of neural markers (*Sox1, Sox2* and *Hes5*) with a concomitant shift of its NMP index to neural and a loss of mesodermal potential. Furthermore, inhibition of Wnt in the Epi-meso2 state (EM2-FP, Fig. 3b-c and Supplementary Fig. S6b-c) leads to a reduction in the expression of neural progenitor markers and an elevation in the expression of mesodermal ones (*Gata6* and *Snail1*), which appropriately shift its NMP index to the mesodermal side with low neural potential in comparison to Epi-meso2.

### Epi-NMPs and the Epi-meso contribute to axial extension

During the elongation of the posterior body axis, NMP derivatives undergo a progressive exit from their niche adjacent to the node and enter either the primitive streak, where they ingress and integrate with the presomitic mesoderm, or are retained in the epiblast and enter the posterior region of the spinal cord. Depending on the time of exit of the niche, they will contribute to different anteroposterior axial levels. With this in mind, we wanted to test the ability of the in vitro derived cells to exhibit these behaviours when transplanted in vivo. To follow the behaviour of transplanted NMP-like populations via live imaging and test the developmental potential these cells have, we decided to use chicken embryos, as these embryos have been shown to be reliable hosts for the functionality of neural and mesodermal progenitors (Fontaine-Perus et al., 1997; Fontaine-Perus et al., 1995; Gouti et al., 2014). Application of this technique to ESCs and different ESCs derived NMPs-like, confirms that it is a reliable assay of developmental potential (Baillie Johnson et al., 2018; Gouti et al., 2014).

We decided to focus our experiments on the EpiSCs derived NMP-like populations as the different tests suggest that they are the closest to the embryo NMPs. In our experiments we transplanted cells from EpiScs, Epi-NMPs and Epi-meso conditions. As can be seen in Fig. 5c EpiSCs contribute only to short axial extensions and their descendants mainly located in the mesodermal region. This result suggests that the EpiSCs exit early from the NMP domain. On the other hand, Epi-NMPs and Epi-meso sequentially contribute to more posterior regions (Fig. 5c) with both neural and mesodermal contributions; we notice that the Epi-meso make more dual neural and mesodermal contributions than the Epi-NMPs and show a weak, but noticeable, bias towards more posterior positions. Since the Epi-meso population is derived from Epi-NMPs, these results suggest that their temporal sequence in vitro results in cells with more posterior colonizing ability in the transplants. Perhaps this reflects the fact that Epi-meso cells express more posterior Hox genes than Epi-NMPs (Fig. 2b and Fig. S4) and this might contribute to their ability to colonize more posterior regions of the embryo (see (Denans et al., 2015))

**Fig 5.**
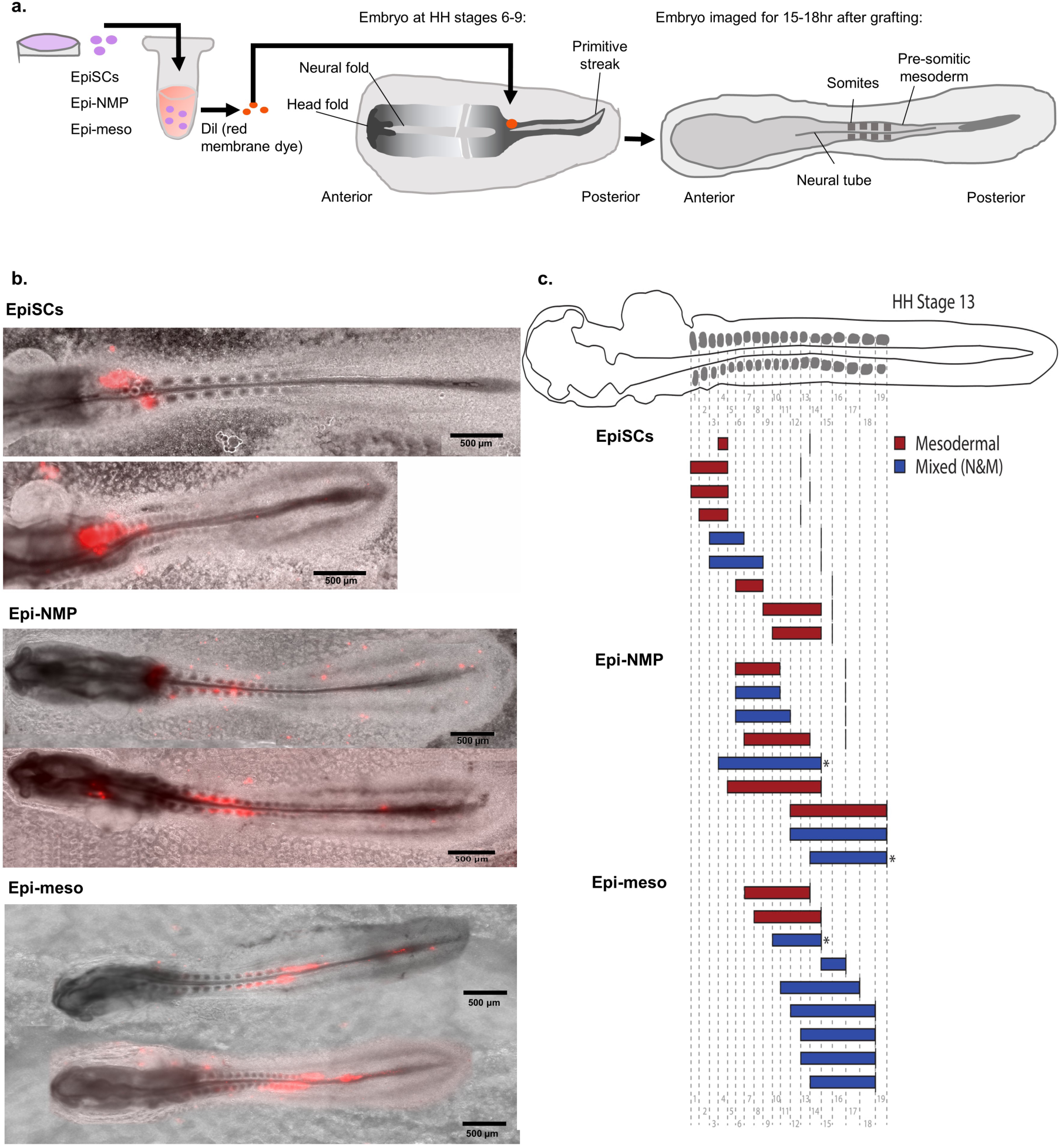
Epi-NMP and Epi-meso progressively contribute to more posterior portions of the embryonic body axis. **a**. Scheme of grafting cells into chicken embryos: colonies from EpiSCs, Epi-NMP and Epi-meso were labelled with membrane dye (Dil; see Methods). The labelled tissues were grafted into the region of the caudal lateral epiblast of a chicken embryo around HH stages 6-9. Embryo cultures were imaged as single time points and as time lapses for 15–18 hours after grafting (see Methods). **b**. Representative images of labelled EpiSCs, Epi-NMP and Epi-meso grafts (marked in red) after transplantation. The red cells represent the contribution of the EpiSCs/Epi-NMP/Epi-meso grafts in the chicken embryo after 15-18hr since grafting. **c**. Schematic of graft contributions in HH stage 13. The length of each labelled cell contribution is shown as a solid bar, coloured red where the cells were found in mesodermal compartments only, or blue where the cells additionally contributed to neural tissue. The length of each contribution is represented as the level of the somites in which labelled cells were found; a solid line denotes the caudal boundary of the most recently formed somite in each case. Where the bar abuts the line, labelled cells could be found into the unsegmented region of the body axis. Contributions greater than 1750μm in length are denoted with asterisks.

Taking the above result together with our gene expression analyses, we conclude that the continued propagation of Epi-NMP population in culture can lead to the production of a population that closely resembles the arisen E8.25 embryonic NMPs.

## Discussion

We have used and compared three PSCs based differentiation protocols to study the emergence in vitro of a population of bipotential progenitors, NMPs, that in the mammalian embryo, give rise to the paraxial mesoderm and spinal cord of the thoracic tract. Our results show that each of these protocols produces populations of cells with different gene expression signatures and ability to contribute to axial elongation but with two common denominators: coexpression of *T* and *Sox2* as well as of genes associated with LPM, IM and allantois. These results suggest that coexpression of *T* and *Sox2* is not a univocal criterion to identify NMPs, that the populations generated in vitro are not restricted to NMPs and that, therefore, the identification of these progenitors requires additional criteria, in particular an ability to self renew and to make long contributions to axial extension as well as an association with the node (Gouti et al., 2015; Henrique et al., 2015; Steventon and Martinez Arias, 2017; Wilson et al., 2009). Applying these criteria to differentiating PSCs populations, we identify a specific protocol that starting with EpiSCs yields a population similar to the NMPs in the embryo in terms of cellular function, gene expression, self renewal and the exit timing of the progenitors from the caudal domain of the embryo. This population emerges from a late epiblast like state that can also give rise to LPM, IM and extraembryonic mesoderm in a signalling dependent manner.

Our observations suggest that a multipotent population might be an obligatory intermediate for the emergence of the NMPs. ESC based protocols yield similar populations that can be differentiated into mesodermal and neural progenitors but lack several features characteristic of NMPs, in particular their ability to self renew and to contribute significantly to axial extension (Baillie Johnson et al., 2018; Gouti et al., 2014; Turner et al., 2014). Furthermore, as we have shown here, these populations represent highly heterogeneous populations with a low representation of NMPs

There are many studies in which Wnt signalling can caudalize epiblast like populations (Amin et al., 2016; Mazzoni et al., 2013; Neijts et al., 2016; Nordstrom et al., 2002; Nordstrom et al., 2006) and in these cases, which are mostly ESCs based, the NMP-like cells fail to self renew as they do in vivo. In contrast to these ESCs based protocols, here we have shown that exposure of pre-treated EpiSCs to FGF and Chiron generates a population with a gene expression signature characteristic of a late CE, around the time of the appearance of the node. Upon further exposure to Wnt and FGF signalling, this population evolves and generates cells with many of the hallmarks of the NMPs, including limited but robust self renewal, as well as an ability to differentiate into neural and mesodermal progenitors in a Wnt dependent manner and make long and more posterior contributions to axial extension in a xenotransplant assay. For these reasons, we propose to call the Epi-NMP population Epi-CE, and the Epi-meso, Epi-NMP.

A significant feature of the Epi-CE cells, in common with the ESCs derived populations, is the expression of markers for LPM, IM and allantois, suggesting that, in vitro, the NMPs are derived from a multipotent population that is likely to exist also in the embryo. Analysis of lineage tracing data at the single cell level supports the existence of this population in the form of clones that span the spinal cord and, at least, two mesodermal derivatives (see Figure 4 in (Tzouanacou et al., 2009)). We find that the fate of the Epi-CE cells is dependent on a balance between BMP and Nodal signalling and has a strict requirement for Wnt signalling in both neural and mesodermal lineages. The dependence on BMP and Nodal mirrors events in the embryo where BMP signalling is concentrated at the caudal end and promotes posterior (LPM, IM and allantois (Sharma et al., 2017)) fates whereas Nodal, expressed around the node, promotes anterior (NMPs) fates.

The importance of Nodal in the establishment of the multipotent population, and perhaps also in the definition of the NMP domain, is underscored by our studies with Nodal mutant cells in which the rescue of a population with disrupted relative level between *Sox2* and *T*, is crucially dependent on the levels of Nodal signalling. Consistent with a role of the node and of Nodal in this population, embryos mutant for *Foxa2* which lack a node, exhibit deficiencies in the organization of the CE and axial elongation (Ang and Rossant, 1994; Weinstein et al., 1994) and the same can be observed in embryos mutant for *Smad2* and *Smad3* (Vincent et al., 2003).

In vitro, the transition between the Epi-CE and Epi-NMP is linked to the loss of expression of several genes that are associated with the epiblast e.g. *Fgf5, Otx2* and, specially, *Oct4*, a POU domain transcription factor that, together with *Sox2*, maintains pluripotency. A similar transition can be observed in the embryo where *Oct4* expression ceases at around E8.5/9.0 (Downs, 2008; Osorno et al., 2012), the time at which cells start differentiating. It is possible that the combination of *Oct4* and *Sox2* promotes multipotency and that only when *Oct4* expression ceases then *Sox2* starts playing a proneural role. A function for *Oct4* in axial elongation can be gauged from the severe axial truncations that follow loss of *Oct4* activity from E7.0/7.5 (DeVeale et al., 2013) and the extended axial elongations associate with overexpression of *Oct4* (Aires et al., 2016). This may reflect an increase in the initial size of the multipotent CE pool rather than a specific alteration in the NMP population.

During the passage of the Epi-NMP population in the presence of Wnt and FGF signalling, we notice that cells progressively lose *T* expression and increase *Sox2* expression. This is surprising since a widespread notion suggests that Wnt signalling suppresses neural differentiation and promotes mesoderm. However, while in the embryo this is true during the first phase of gastrulation, before the appearance of the node at E7.5, this might not be the case during the development of the caudal region of the embryo. In fact, during gastrulation, Wnt signalling does not suppress neural but rather anterior epiblast which is committed to anterior neural (Arkell and Tam, 2012; Lewis et al., 2008). When the trunk develops, there is a clear evidence that Wnt/β-catenin signalling is required for the expansion of the neural progenitors in the spinal cord (Zechner et al., 2003) and an increase in Wnt/β-catenin signalling does not supress neural development (Garriock et al., 2015). Thus, we suggest that the response of neural specified cells to Wnt signalling is a measure of the stage and position of the cells. A requirement for Wnt signalling in the development of the spinal cord is further emphasized by the observation that the *Sox2* gene has a Tcf response element and responds to Wnt signalling (Takemoto et al., 2006).

In summary, using a specific experimental protocol we have shed light on the origin of the NMP population in vivo and in vitro. Significantly the Epi-NMP population exhibits a limited but robust self renewing ability which, together with its gene expression signature, lead us to believe that there is an in vitro correlation with the in vivo NMPs.

## Acknowledgements

This work was supported by Cambridge Trust and Cambridge Philosophical Society scholarships and AJA Karten trust award to S. Edri, a Sir Henry Dale Fellowship jointly funded by the Wellcome Trust and the Royal Society (GrantNumber 109408/Z/15/Z) to B. Steventon, an EPSRC studentship to P.Baillie-Johnson and BBSRC project grants (No. BB/M023370/1 and BB/P003184/1) to AMA. We are grateful to J Collignon for the Nodal mutant cells and to James Briscoe, Meritxell Vinyoles, Vikas Trivedi and Valerie Wilson for discussions.

## Materials and Methods

### Cell culture

E14-Tg2A were grown in tissue-culture plastic flasks coated with 0.1% gelatine (Sigma-Aldrich, G1890-100G) in PBS (with Calcium and Magnesium, Sigma-Aldrich, D8662) filled with GMEM (Gibco, UK) supplemented with non-essential amino acids, sodium pyruvate, GlutaMAX™, β-mercaptoethanol, foetal bovine serum and LIF. Cell medium was changed daily and cells passaged every other day. The differentiation protocols are as the following:

#### ES-NMP

Cells were plated at a density of 4.44×10^3^ cells/cm^2^ in a 0.1% gelatine coated flask with a base medium of N2B27 (NDiff 227, Takara Bio) for 2 days. After 48hr N2B27 is supplemented with 3μM of CHIR99021 (Chiron 10mM, Tocris Biosciences) for additional 24hr, which are in total 72hr.

#### ES-meso and ES-neuro

ES-NMP cells were detached from the culture flask using Accutase (BioLegend 0.5Mm) and divided into 2 flasks coated with 0.5% Fibronectin at a dense of 7.5×10^3^ cells/cm^2^. To get ES-neuro the cells were grown for 2 days in N2B27 and for ES-meso the cells were grown in N2B27 supplemented with 20ng/ml FGF2 (R&D systems, 50μg/ml) and 3μM Chiron.

#### ES-NMPF, ES-neuroF, ES-mesoF (Gouti et al., 2014)

Cells were plated at a density of 5×10^3^ cells/cm^2^ in a 0.1% gelatine coated CellBINDSurface dish (Corning) with a base medium of N2B27 supplemented with 10 ng/ml FGF2. After 48hr N2B27 is supplemented with 10 ng/ml FGF2 and 5μM Chiron for additional 24hr, which are in total 72hr. To induce neural SC identity (ES-neuroF) or mesodermal identity (ES-mesoF) day 3 – day 5 was followed by either N2B27 supplemented with 100nM RA (Sigma) or by N2B27 supplemented with 5μM Chiron respectively.

#### Epi-NMP

E14-Tg2A were grown in tissue-culture plastic flasks coated with 0.5% Plasma Fibronectin (FCOLO, 1mg/ml, Temecula) in PBS (with Calcium and Magnesium) filled with N2B27 supplemented with 12ng/ml FGF2 and 25ng/ml Activin A (Stem Cells Institute 100μg/ml), this known as Epi-media with or without 20μM XAV939 (XAV Tocris Biosciences, 10mM) for at least 4 passages. Those cells known as EpiSCs (or EpiXAV when the β-catenin inhibitor XAV is used). Those cells can be tested to be EpiSC by seeding them in a colony assay density (67 cells/cm^2^) in restricted medium (2i: N2B27 supplemented with 3μM Chiron and 1μM PD0325901 (PD03, Tocris Biosciences, 10mM)), resulting in no growth of cells, ensuring that the cells are no longer in the naïve pluripotent state and they moved on to the prime pluripotent state (data are not shown).

EpiSCs (treated with or without XAV) were plated at a 5×10^4^ cells/cm^2^ density in a 0.5% Fibronectin pre-coated flask with Epi-media for the first day. Day 2 is followed by increasing the concentration of FGF2 to 20ng/ml in the base medium of N2B27 and removing Activin A or XAV if originally was used to grow the EpiSCs. On day 3, N2B27 is supplemented with 3μM Chiron which is added to the 20ng/ml FGF2. After 72hr those cells known as the Epi-NMP or EpiXAV-NMP if XAV was used in the Epi-media. This protocol is a variation of one that has been used to derive NMP-like cells from human ESCs (Lippmann et al., 2015)

#### Epi-meso and Epi-neuro

Epi-NMP (without XAV) cells were detached from the culture flask using Accutase and divided into 2 flasks coated with 0.5% Fibronectin at a dense of 5×10^4^ cells/cm^2^. To get Epi-neuro the cells were grown for 2 days in N2B27 and for Epi-meso the cells were grown in N2B27 supplemented with 20ng/ml FGF2 and 3μM Chiron.

#### Epi-meso differentiation

Epi-meso (without XAV) cells were detached from the culture flask using Accutase and plated back to a 0.5% Fibronectin coated flask at a dense of 5×10^4^ cells/cm^2^ for 2 days in N2B27 supplemented with 20ng/ml FGF2 and 3μM Chiron. The first passage from Epi-meso is named Epi-meso2 (EM2), the second passage is named Epi-meso3 (EM3) and so forth.

#### BMP, FGF and Wnt signalling

##### Epi-mesoFCD

Epi-NMP (without XAV) cells were detached from the culture flask using Accutase and plated back to a 0.5% Fibronectin coated flask at a dense of 5×10^4^ cells/cm^2^ for 2 days in N2B27 supplemented with 20ng/ml FGF2, 3μM Chiron and 500nM dorsomorphin-H1 (DMH-1 5mM, Tocris Biosciences) which is a BMP inhibitor.

##### EM2-FCD

Epi-meso (without XAV) cells were detached from the culture flask using Accutase and plated back to a 0.5% Fibronectin coated flask at a dense of 5×10^4^ cells/cm^2^ for 2 days in N2B27 supplemented with 20ng/ml FGF2, 3μM Chiron and 500nM DMH-1.

##### EM2-FCB

Epi-meso (without XAV) cells were detached from the culture flask using Accutase and plated back to a 0.5% Fibronectin coated flask at a dense of 5×10^4^ cells/cm^2^ for 2 days in N2B27 supplemented with 20ng/ml FGF2, 3μM Chiron and 1ng/ml BMP4 (R&D Systems, 100μg/ml).

##### EM2-Chiron

Epi-meso (without XAV) cells were detached from the culture flask using Accutase and plated back to a 0.5% Fibronectin coated flask at a dense of 5×10^4^ cells/cm^2^ for 2 days in N2B27 supplemented with 3μM Chiron alone.

##### EM2-FP

Epi-meso (without XAV) cells were detached from the culture flask using Accutase and plated back to a 0.5% Fibronectin coated flask at a dense of 5×10^4^ cells/cm^2^ for 2 days in N2B27 supplemented with 20ng/ml FGF2 and 1μM IWP-2 (PIN 5mM, STEMGENT) which is a Wnt pathway inhibitor.

##### Nodal Null cells

Cells mutant for Nodal (Nodal-/-) were provided by J. Collignon and were derived from the 413.d mutant mouse line (Conlon et al., 1991), They were grown on a 0.5% Fibronectin coated culture flask with Epi-media: N2B27 supplemented with 12ng/ml FGF2 and 25ng/ml Activin A for at least 4 passages. The Nodal null EpiSCs were plated in 5×10^4^ cells/cm^2^ density on a 0.5% Fibronectin pre-coated flask with Epi-media for the first day. Day 2 is followed by increasing the concentration of FGF2 to 20ng/ml, in the base medium of N2B27, and removing the Activin A. On day 3, N2B27 is supplemented with 3μM Chiron which is added to the 20ng/ml FGF2. After 72hr those cells known as the Nodal-/- Epi-NMP. In order to examine the role of Nodal in establishing the NMPs, the Nodal mutant Epi-NMPs were supplemented with 2 different doses of Nodal in the culturing medium on the 3^rd^ day: 20ng/ml FGF2, 3μM Chiron and either 100ng/ml of Nodal (R&D systems, sample name: Nodal-/- Epi-NMP+0.1xNodal) or 1 μg/ml of Nodal (sample name: Nodal-/- Epi-NMP+1xNodal) in the base medium of N2B27.

## Quantitative RT-PCR (qRT-PCR)

Total RNA was isolated from cells using Trizol. First strand cDNA synthesis was performed with Superscript III system (Invitrogen) and the quantification of doublestranded DNA obtained with specific genes designed primers, using QuantiFast SYBR Green PCR Master Mix (Qiagen) and the standard cycler program (Qiagen RotorGene Q). The qPCR was done in technical triplicates. The primers that have been used are available in Supplementary Material Table S1. All experiments were performed in biological duplicates or triplicates. Expression values were normalized against the housekeeping gene *Ppia*. To compare different experiments of qRT-PCR, each run of the qPCR included one common condition, in our case it was the Epi-meso. Each condition in every run was normalized to Epi-meso and averaged across biological replicas. Here are the steps to calculate the normalized gene expression values:

1. Identifying the C_t_ (threshold cycle) for each gene (technical triplicates) and calculating the expression values (2^−Ct^).
2. Calculating the average and the standard deviation (std) for each gene from the triplicate expression values.
3. Dividing the average and the std of each gene in the expression value of *Ppia*.
4. The normalized gene expression values in condition x will be divided with the normalized gene expression values in the common condition in every qRT-PCR experiment (Epi-meso) as the following: F=x/y, where x denotes the expression of a gene at any condition and y denotes the expression of the same gene at Epi-meso condition. Both x and y have an error, the std that have been calculated in step 3: Dx and Dy, hence the total error is:

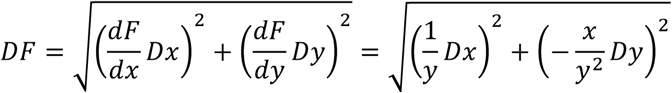
5. Averaging the biological replicas: F_1_ and F_2_ are biological replicas of the same gene in the same condition and their expression was normalized as the above steps. The average of the normalized expression and the error is calculated as the standard error:

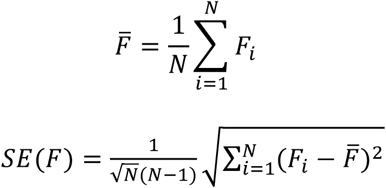

where N is the number of biological replicas (between 2 and 3).
6. Standardizing the normalized expression values of a gene to Z-score values across conditions was done according the equation below:

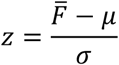 Where μ and σ denote the average and the standard deviation of the normalized expression of a gene across all the conditions examined in this work respectively (the average and standard deviation of 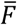 for a specific gene across all the conditions).

## NMP and Epiblast indices

| Neural Genes | Mesodermal genes | Epiblast genes |
| --- | --- | --- |
| *Sox1, Pax6, Hes5, Sox2* | *Bmp4, Evx1, Gata6, Meox1, Mesp1, Mixl1, Raldh2, Tbx2, Tbx6, Msx1, Msx1, Pax2, Osr1, Snai1, T* | *Cdh1, Fgf5, Oct4, Otx2, Cdx2, Fgf8, Nodal, Wnt3a, Cyp26a1, Nkx1-2, Hoxc6, Cdh2, Foxa2* |

The NMP index was calculated as the following: In all the conditions (17 in total) the average expression Z-score value of the neural genes and the mesodermal genes (noted in the table above) was obtained and scaled between 0 – 1 across the 17 conditions. This resulted in 2 values for each condition: the neural potential and the mesodermal potential. The epiblast index was calculated in a similar manner: the average of the Z-score expression values of the epiblast genes was calculated versus the differentiation genes (neural and mesodermal, listed in the table above) and scaled between 0 – 1 across the 17 conditions, resulting in an epiblast potential and a differentiation potential for each condition.

### Single molecule fluorescence *in-situ* hybridization (sm-FISH)

Single molecule RNA FISH was carried out as described previously (Nair et al., 2015). Cells were dissociated using Accutase, washed in PBS, fixed in 37% formaldehyde at room temperature and permeabilized and stored in 70% ethanol at 4°C. All washes and hybridizations were carried out in suspension. Wash buffers included 0.1% Triton X-100 to minimize losses of cells sticking on the walls. Samples were mounted between a slide and #1 cover glass, in the glucose-oxidase-based 2 × SSC anti-fade buffer. The probes for the genes (supplementary Table S2) were designed using StellarisTM website and bought via StellarisTM FISH probes (Biosearch Technologies) (Raj et al., 2008). Additional information about how the probes were designed, prepared and used can be found in (Raj et al., 2008). Cells were imaged within 24 to 48h on a Nikon Ti-E wide field microscope, using a 60X oil-immersion objective and a cooled camera (Orca flash 4.0, Hamamatsu). The cells in the images were segmented manually and the spot-detection was done semi-automatic using a MATLAB graphic user interface (GUI) developed by Marshall J. Levesque and Arjun Raj at the University of Pennsylvania or home-made protocols written in ICY (de Chaumont et al., 2012).

### Principal Component Analysis

PCA (Nair et al., 2015) involves the assignment of data, in our case gene expression, to new coordinates named principal components or PCs. The variance of observed coordinates in each PC occurs in a decreasing order, observations (the samples) projected on PC1 have a greater variance than the same observations projected on PC2 and so on. The PCs were calculated according to the Z-scores expression values of the 27 genes expressed (see Fig. 2b and Fig. S4b) in the different stages of the mouse embryo epiblast/ectoderm and in the 3 in vitro protocols and their neural and mesodermal differentiation: ES-NMP, ES-NMPF and Epi-NMP.

### Cosine similarity

Here we used cosine similarity as a measure of similarity between Z-scores expression values of list of genes in one condition versus other condition (i.e. Epi-NMP versus the mouse embryo epiblast stages (Kojima et al., 2014) per the same list of genes). The cosine similarity was calculated as the following:

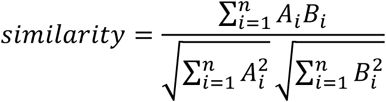

Where A and B represents the list of genes with their values of Z-score gene expression in two conditions and A_i_ and B_i_ are the components of these two vectors. The similarity was constrained to the positive space, where 0 indicates that the two vectors are dramatically opposite and 1 indicates maximal similarity.

### Confocal and immunostaining

In 4-well (Ibidi), plastic tissue-culture dishes the different samples were grown. Samples were washed in BBS + CaCl_2_ (50 mM BES Sodium Salt, 280 mM NaCl, 1.5 mM Na2HPO4, 1 mM CaCl_2_ adjusted to pH 6.96 with 1 M HCl) and fixed for 15 minutes in 4% paraformaldehyde. Samples were washed and permeabilised with BBT (BBS + CaCl_2_ supplemented with 0.5% BSA and 0.5% Triton X-100) before overnight antibody staining, following a wash with BBT and then the samples were incubated for 2hr with the desired fluorescently-conjugated secondary antibody. Prior to imaging, samples were washed with BBS + CaCl_2_ and covered in a mounting medium (80% spectrophotometric grade glycerol, 4% w/v n-propyl-gallatein in BBS + CaCl_2_).

The following primary antibodies were used: *T (Brachyury)* N19 (goat; Santa Cruz Biotechnologies, sc17743, dilution 1:100), *Oct3/4* (mouse; Santa Cruz Biotechnologies, sc5279, dilution 1:100), *Sox2* (rabbit; Millipore, AB5603, dilution 1:200), *Otx2* (goat, R&D AF1979 dilution 1:200), *Cdh2* (mouse, BD Bioscience 610920 dilution 1:200) and *Cdh1* (rat, Takara M108, dilution 1:100). Secondary antibodies (Goat-A488, Rabbit-A633, Mouse-A568, Rat-A633) from Molecular Probes, were made in donkey and used in a 1 in 500 dilution with Hoechst 33342 (H3570, dilution 1 in 1000; Invitrogen ThermoFisher). Samples were imaged using an LSM700 on a Zeiss Axiovert 200 M with a 63× EC Plan-NeoFluar 1.3 NA DIC oil-immersion objective. Hoechst, Alexa488, -568 and -633 were sequentially excited with a 405, 488, 555 and 639 nm diode lasers, respectively. Data capture carried out using Zen2010 v6 (Zeiss).

### Chicken Embryo Culture

Fertilised chicken eggs were stored in a humidified 10°C incubator for up to one week until required. Eggs were transferred to a humidified, rocking 37°C incubator for 24 hours prior to the preparation of embryo cultures, which was done according to a modified version of New Culture (New, 1955). Embryo cultures were incubated at 37°C prior to grafting and were fixed in 4% paraformaldehyde within 24 hours.

*Graft Preparation and Transplantation:* Cell cultures were prepared as described above. Adherent cell cultures were detached mechanically using a cell scraper in PBS (with calcium and magnesium) to lift intact colonies with minimal sample dissociation. The tissues were labelled by transferring them to a FBS-precoated FACS tube and were centrifuged at 170 × *g* for five minutes. The supernatant was discarded and the colonies washed by gentle resuspension in PBS (with calcium and magnesium), before the centrifugation step was repeated. The tissues were then resuspended gently in PBS (without calcium and magnesium) for labelling with DiI (Thermo Fisher Scientific Vybrant^®^ V22885, 1% v/v) for 25 minutes in the dark, on ice. The labelled tissues were centrifuged at 170 × *g* for five minutes and the pellet was gently resuspended in PBS (with calcium and magnesium) for grafting.

Labelled tissues were grafted into the region of the chick caudal lateral epiblast as described in (Baillie Johnson et al., 2018; Gouti et al., 2014), using an eyebrow knife tool. Embryo cultures were imaged as single time points and as time-lapses for 15–18 hours after grafting.

### Embryo Microscopy and Image Analysis

Widefield images were acquired with a Zeiss AxioObserver Z1 (Carl Zeiss, UK) using a 5x objective in a humidified 37°C incubator, with the embryo cultures positioned on the lid of a six-well plate. An LED white light illumination system (Laser2000, Kettering, UK) and a Filter Set 45 filter cube (Carl Zeiss, UK) was used to visualise red fluorescence. Emitted light was recorded using a back-illuminated iXon800 Ultra EMCCD (Andor, UK) and the open source Micro-Manager software (Vale Lab, UCSF, USA). The open source FIJI ImageJ platform (Schindelin et al., 2012) and the pairwise stitching plugin (Preibisch et al., 2009) were used for image reconstruction and analysis.

